# Cell sorting based on single nucleotide variation enables characterization of mutation-dependent transcriptome and chromatin states

**DOI:** 10.1101/2024.07.05.602247

**Authors:** Roberto Salatino, Marianna Franco, Arantxa Romero-Toledo, Yi Wang, Shanel Tsuda, Oszkar Szentirmai, Michalina Janiszewska

**Affiliations:** Department of Molecular Medicine, The Herbert Wertheim UF Scripps Institute for Biomedical Innovation & Technology, Jupiter, FL, USA; Department of Immunology and Microbiology, The Herbert Wertheim UF Scripps Institute for Biomedical Innovation & Technology, Jupiter, FL, USA; The Skaggs Graduate School of Chemical and Biological Sciences, The Scripps Research Institute, La Jolla, California, USA; Center for Neurological Surgery and Neurosciences, Cleveland Clinic Martin Health, Port St. Lucie, Florida USA

**Keywords:** STAR-FACS, SNV, single cell, cell sorting, TERT promoter mutation, CUT&Tag

## Abstract

Single nucleotide variants (SNVs) contribute to cancer by altering the coding and the non-coding regions of the genome. Connecting SNVs to transcriptomic and epigenetic changes at the single-cell level remains challenging. To enable studies of rare cell populations harboring specific point mutations, we developed STAR-FACS, Specific-To-Allele PCR-FACS, to sort cells based on genomic allele alterations. We show that STAR-FACS can separate cells based on TERT promoter mutation status and is compatible with bulk and single-cell transcriptomic and epigenetic profiling. We demonstrate that glioblastoma cell lines derived from the same tumor but harboring distinct TERT promoter SNVs have different transcriptional programs.

## BACKGROUND

Hotspot mutations, occurring at specific locations within oncogenes and tumor suppressor genes, are considered to be the major genetic drivers of tumorigenesis[1,2]. These single nucleotide substitutions are recurrent among cancer patients, pointing to strong evolutionary pressure that selects for cancer cells with these alterations[2]. This is due to single nucleotide variation (SNV) resulting in gain or loss of function of the affected gene. In addition, hotspot mutations can also have profound effects on non-coding regions of the genome, by creating novel transcription factor binding sites and affecting chromatin structure[3]. Whole exome and whole genome sequencing of cancer tissues identified multiple cancer driver mutations in coding and non-coding portions of the genome[2,3] and linked these SNVs to the development of therapy resistance[4].

Many of the SNVs of clinical significance, linked to therapy resistance, are non-clonal in untreated tumors and only present in a subpopulation of cells within the tumor tissue[5–7]. Intratumor heterogeneity, linked with poor prognosis across multiple cancer types, can be assessed using SNVs to infer the clonal architecture of the tumor[5,7–9]. Nonetheless, bulk sequencing methods are limited in their sensitivity to detect rare SNV-harboring subpopulations, which makes it challenging to study the functional effects of subclonal mutations. Since clonal interactions are prevalent in heterogenous tumors[10–12], as well as in normal tissue with rare mutant cells[13,14], studying the phenotypes driven by hotspot mutations in cells experiencing competition or cooperation is of high interest. Single-cell whole genome and whole exome sequencing could address these interactions[15–19], although high cost of these experiments and high error rates hinder the wide adoption of these techniques. Novel technologies are now enabling the detection of predefined panels of SNVs at the single-cell level, increasing the signal-to-noise ratio[20,21]. Single-cell co-sequencing of DNA and RNA from individual nuclei could overcome the challenges of direct measurement of a transcriptomic or epigenomic feature associated with a specific genomic alteration [22]. However, all of these techniques require specialized equipment and consumables, making them costly and not easy to scale for a large number of cells.

Here, we introduce a new method for fluorescence-based sorting and enrichment of cells based on their genomic SNV status. In the Specific-To-Allele PCR FACS (STAR-FACS) protocol, paraformaldehyde-fixed intact cells are subjected to *in cell* PCR amplification to generate a mutation-specific amplicon which can be labeled with fluorescent probes, enabling FACS sorting of the mutant cell population. Our novel mutation-based cell enrichment method is inexpensive and compatible with bulk and single-cell transcriptomic characterization of the mutant and wild-type populations, as well as with chromatin state profiling by CUT&Tag. The application of STAR-FACS to glioblastoma (GBM) primary cell lines reveals transcriptional divergence between TERT promoter mutant and wild-type subclonal populations derived from different regions of the same tumor.

## RESULTS

### Overview of STAR-FACS

We adapted our previously developed method for in situ mutation detection at the single-cell level, STAR-FISH[23–25], to enable intact cell sorting based on the presence or absence of a mutation in their genomic DNA. The assay development was focused on the detection of the most common cancer mutation in a non-coding region, namely the telomerase reverse transcriptase promoter (TERTp) mutation[3]. Two hotspot mutations, C228T and C250T (C to T transitions at 124 and 146 base pairs, respectively, upstream of TERT ATG start codon), account for 80-90% of mutations in TERTp[3]. Each of these variants results in a novel binding site for the GAPB transcription factor, which activates telomerase expression, enabling telomere extension, escape from replicative senescence and cell immortalization[26]. In addition, TERTp mutations disrupt the G-quadruplex structure of the promoter, resulting in long-range changes in chromatin organization[27,28]. In glioblastoma, the hotspot mutations are present in over 80% of tumors, however they are not always clonal[29,30]. Detection of a TERTp mutation and its resulting TERT reactivation is challenging with traditional expression-based methods, due to the relatively low abundance of the transcript and protein.

To enable studies of transcriptomic and epigenetic changes driven by subclonal TERTp mutations, we first established a PCR protocol to selectively generate an amplicon from TERTp C228T and C250T mutant loci. Two model glioblastoma cell lines known to harbor these mutations were used as a model system, U87-MG (C228T mutant) and DK-MG (C250T mutant). We modified the STAR-FISH protocol to enable *in cell* PCR (**Fig. 1** and **Methods**). Briefly, single cell suspension was fixed, permeabilized and frozen in 5% DMSO in PBS on Day 1. On Day 2, two rounds of *in cell* PCR with OneTaq Hot Start polymerase were performed. The 1^st^ round of PCR was used to amplify the 597bp region of the promoter containing both hotspots and the 2^nd^ round included mutation-specific primers containing unique 20bp overhangs not found in the human genome. After washing, the cells were subjected to a DNA hybridization step with fluorescently labeled probes complementary to the overhangs introduced in the 2^nd^ round of *in cell* PCR. High stringency washes were implemented to ensure the removal of unhybridized probes. After these steps, cells retain their intact morphology and can be directly sorted based on the signal from the fluorescently labeled amplicon.

**Figure 1.**
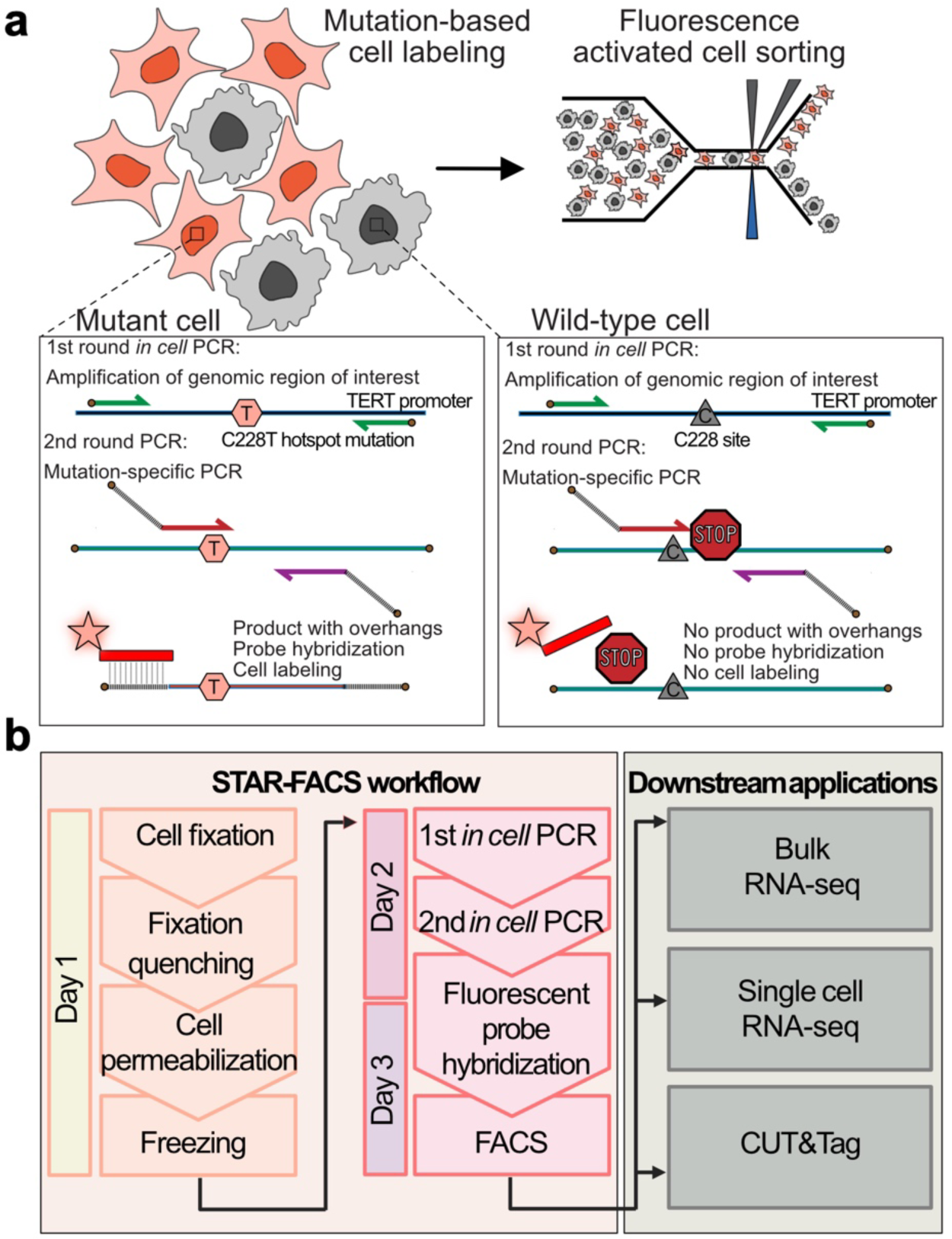
STAR-FACS protocol outline. **a**) Principle of mutation-based cell labeling with STAR-FACS for TERT promoter hotspot mutation C228T. First, the genomic DNA region of interest (597bp fragment of TERT promoter) in intact fixed cells is subject to PCR amplification. In the second round PCR, mutation-specific primers with unique overhangs are used. The primers contain mismatches, enabling efficient amplification of the mutant, but not the wild-type amplicon. The unique overhangs incorporated into the PCR product in the 2nd round PCR allows for specific hybridization of fluorescently labeled probes. Labeled intact cells can be then FACS-sorted. **b**) Fixed and permeabilized cells can be cryopreserved and labeled by two-step *in cell* PCR followed by fluorescent probe hybridization. This enables the sorting of cells for downstream applications, including RNA-seq, scRNA-seq and CUT&Tag.

### TERTp mutation-based cell labeling and transcriptomic analysis

First, we verified that U87-MG cells, harboring C228T mutation, can be fully labeled with the STAR-FACS protocol (**Fig. 2a**). Mixing equal number of U87-MG with DK-MG cells, which harbor the C250T mutation, and labeling them with the C228T-specific STAR-FACS showed expected 1:1 ratio of the labeled to unlabeled cells (**Fig. 2a, Supplementary Fig. 1a, b**). Next, we tested whether combining STAR-FACS for the two TERTp hotspots would retain its specificity. In this experiment, primers specific for C228T and C250T were mixed in a single reaction. Since the primers specific for the two mutations were designed to have distinct overhangs, the two amplicons can be distinguished by hybridization of distinct probes, with an Alexa-647-labeled probe for C228T and a Cy3-labeled probe for C250T. Indeed, a range of ratios of mixed U87-MG/DK-MG cell suspensions can be deconvoluted by STAR-FACS (**Fig. 2b, Supplementary Fig. 1c, d**).

**Figure 2.**
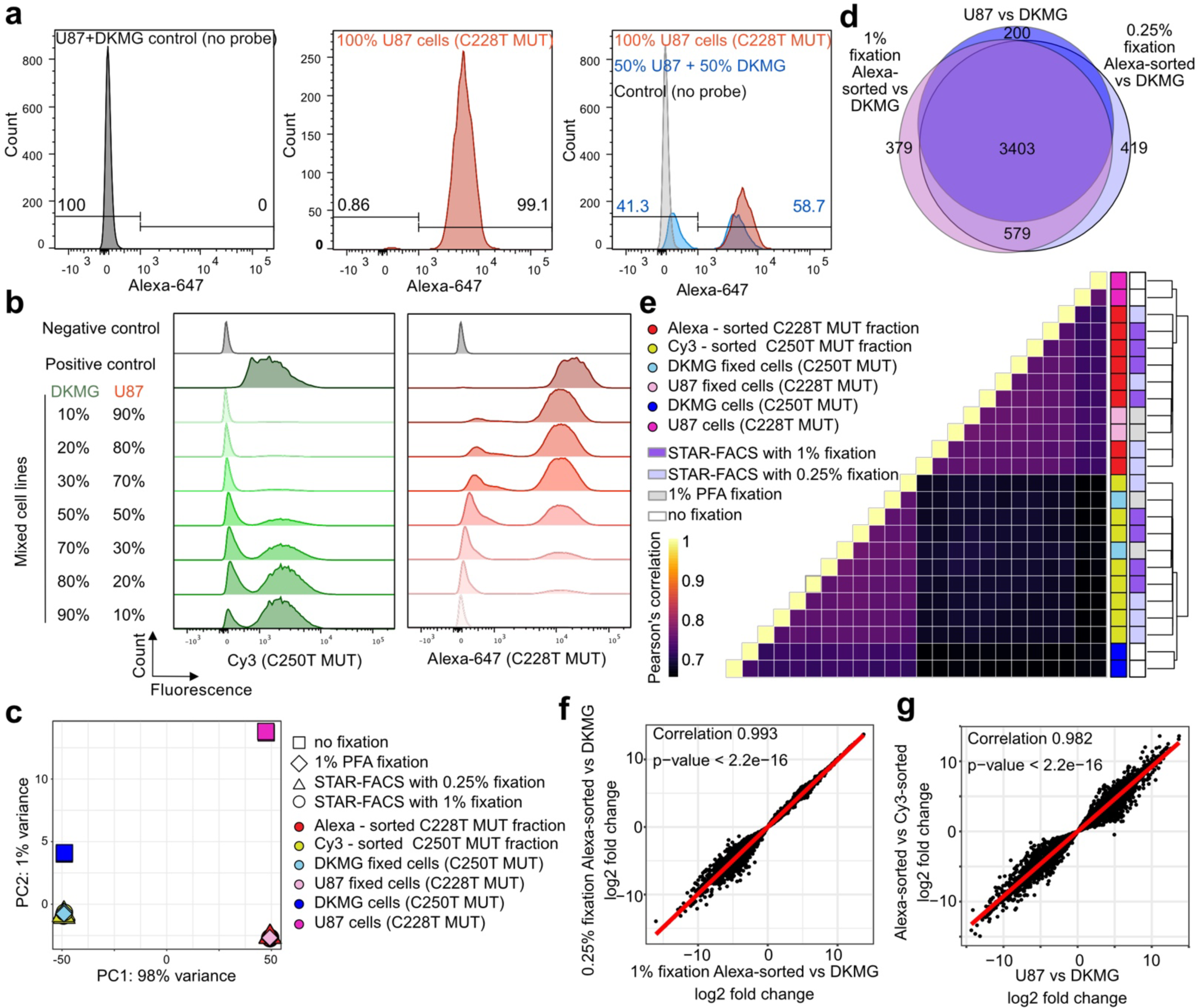
STAR-FACS validation. **a**) STAR-FACS labeling of C228T mutant cells in a mix of U87-MG and DK-MG cells. Left panel – no probe STAR-FACS control. Middle panel – 100% U87-MG cells. Right panel – an overlay of positive (100% U87-MG) and negative control (no probe, 50% U87-MG + 50% DKMG) over STAR-FACS labeled 1:1 mix of U87-MG and DKMG cells. **b**) STAR-FACS labeling for C250T (Cy3-labled probe) and C228T (Alexa-647-labeled probe) mutations. U87-MG/DKMG cells mixed at different ratios were used in a single reaction targeting both mutations. **c**) Principal component analysis of STAR-FACS sorted fractions. Comparison with freshly collected cells (no fixation) and cells fixed with 1% paraformaldehyde (PFA). Two different fixation strengths in the STAR-FACS protocol were tested. Two replicates per each sample are plotted. **d**) Overlap of differential gene expression obtained from fixed U87-MG and DKMG cells and STAR-FACS sorted fractions. Alexa-sorted fractions correspond to U87-MG cells. The number of common differentially expressed genes (adjusted p-value < 0.05) is shown in the center. **e**) Pearson’s correlation of gene expression of STAR-FACS labeled and sorted fractions derived from mixed U87-MG and DKMG cells. Each row and column represents a biological replicate. **f**) Correlation of expression of genes differentially expressed between Alexa-sorted fractions obtained from STAR-FACS with 1% or 0.25% fixation and DKMG fixed cells. **g**) Correlation of expression of genes differentially expressed in Alexa-sorted vs Cy-3-sorted STAR-FACS fractions derived from a mix of U87-MG and DKMG cells and direct comparison of gene expression between U87-MG and DKMG fixed cells.

The primary objective of developing STAR-FACS was to enable transcriptomic studies of rare mutant cell populations. We tested a range of fixatives and established that 1% paraformaldehyde (PFA) in PBS or 0.25% PFA in PBS provide sufficient fixation to keep the cells intact during the STAR-FACS protocol (**Supplementary Fig. 2a, b**). Decrosslinking and amplicon removal (see **Methods**) were needed to ensure good quality of RNA extraction (**Supplementary Fig. 2c, d**). Gene expression profiling of U87-MG/DK-MG mixed populations sorted after STAR-FACS for C228T mutation were remarkably similar to profiles obtained from cell lines fixed with 1% PFA and fresh cell lines (**Fig. 2c-g**). The principal component analysis shows that fixation itself, but not STAR-FACS, accounts for the transcriptional divergence from the fresh cells and that fixation strength (1% vs 0.25% PFA) in the STAR-FACS protocol has a negligible effect on transcript diversity when compared with fresh cells fixed with 1% PFA (**Fig. 2c**). Differential gene expression between 1% PFA fixed U87-MG cells and DK-MG cells is captured very well by STAR-FACS sorted samples, with 3,403 differentially expressed genes captured regardless of fixation and STAR-FACS (**Fig. 2d**). Gene expression levels are also highly concordant for samples prepared with different strength of fixation in STAR-FACS (**Fig. 2e, f, Supplementary Fig. 3a-d**). Mutation-based cell sorting generates results that are congruent with the ground truth comparison of fixed U87-MG vs DK-MG cell lines (**Fig. 2g**). This indicates that, despite the harsh PCR and hybridization conditions, STAR-FACS provides high quality cellular material for transcriptomic studies.

Next, we tested our mutation-based cell separation on co-cultures of primary GBM neurosphere cell lines. The cells were mixed in equal ratios and grown for 10-14 days as a co-culture, before labelling by STAR-FACS and RNA sequencing (**Fig. 3a**). We tested an array of cell lines derived from different regions of the same tumor, all wild-type for TERT promoter mutation (OS23-2, OS23-3, OS23-4, OS23-5), each co-cultured with a mutant cell line from a different tumor (OS11-5 or OS25-4). The transcriptomic data principal component analysis shows that all fractions labeled by TERTp-specific STAR-FACS separate well from their TERTp WT co-culture partners (**Fig. 3b, c**). All Alexa-647^-^ fractions derived from distinct regions of OS23 tumor cluster together with their parental monoculture cell lines (**Fig. 3d**), while Alexa-647^+^ fractions cluster with their corresponding TERTp MUT parental cell lines (**Fig. 3d**). These results demonstrate that even a highly complex neurosphere co-cultures can be unmixed using STAR-FACS.

**Figure 3.**
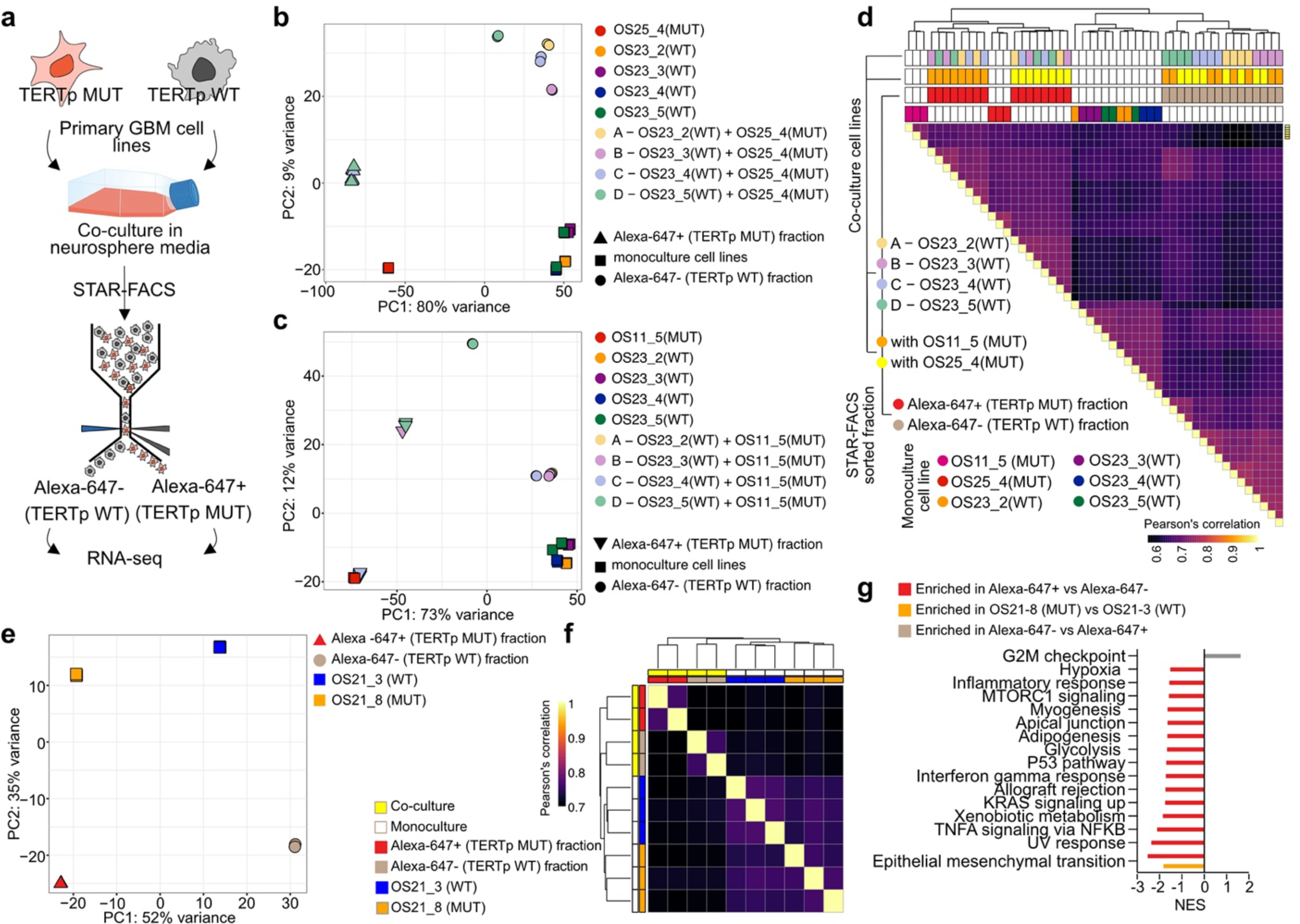
TERT promoter mutation-based cell separation reveals subclonal divergence in GBM. **a**) Outline of the co-culture experiments. TERT promoter mutant (TERTp MUT) cells and TERT promoter wild-type (TERTp WT) cells were mixed in a 1:1 ratio, cultured for 10-14 days and subject to STAR-FACS to separate WT from MUT cells for subsequent transcriptomic analysis. **b**) Principal component analysis of STAR-FACS sorted samples from co-cultures of OS25-4 TERTp MUT cells and WT lines derived from distinct regions of OS23 tumor (OS23-2 – OS23-5). Intense color squares – parental cell lines. Pastel colors – co-cultures. Triangles – Alexa-647+ fractions (TERTp MUT). Circles – Alexa-647-fractions –TERTp WT). Principal component 1 and 2 percent variance is shown. Two biological replicates for each sample are plotted. **c**) Principal component analysis of STAR-FACS sorted samples from co-cultures of OS11-5 TERTp MUT cells and WT lines derived from distinct regions of OS23 tumor (OS23-2 – OS23-5). Intense color squares – parental cell lines. Pastel colors – co-cultures. Triangles – Alexa-647+ fractions (TERTp MUT). Circles – Alexa-647-fractions –TERTp WT). Principal component 1 and 2 percent variance is shown. Two biological replicates for each sample are plotted. **d**) Correlation of gene expression between co-cultured STAR-FACS separated fractions from OS23 tumor. **e**) Principal component analysis of STAR-FACS sorted samples from co-cultures of OS21-8 TERTp MUT cells and WT line (OS21-3), derived from distinct regions of the same tumor (OS21). Squares – parental cell lines. Triangles – Alexa-647+ fractions (TERTp MUT). Circles – Alexa-647-fractions –TERTp WT). Principal component 1 and 2 percent variance is shown. Two biological replicates for each sample are plotted. **f**) Correlation of gene expression between co-cultured STAR-FACS separated fractions from OS21 tumor. **g**) Gene set enrichment analysis of hallmark pathways found when comparing fresh OS21-3 and OS21-8 neurosphere cultures and STAR-FACS fractions. NES – normalized enrichment score. Pathways significantly enriched at nominal p value < 0.05 and FDR < 25% are shown.

Among our cultures derived from distinct regions of the same tumor, we have also identified one tumor with subclonal TERTp mutation, present in the neurosphere culture derived from region 8 (OS21-8), but not in the culture derived from region 3 (OS21-3). Co-culture of these cell lines was also effectively separated by STAR-FACS (**Fig. 3e-g, Supplementary Fig. 3e, f**). Interestingly, STAR-FACS-based cell separation uncovered high expression of G2M checkpoint genes in the TERTp WT fraction, while TERTp MUT fraction was enriched in transcripts playing a role in epithelial to mesenchymal transition (EMT), TNFα/NFκB and KRAS signaling, P53 pathway and glycolysis, hypoxia, and inflammatory responses (**Fig. 3g**). Only EMT was identified as enriched when fresh OS21-8 vs OS21-3 cells were compared. TERT protein binds NFκB to elicit EMT[31] and immune response[32], and has also been linked to glucose metabolism and metabolic reprograming in brain tumors[33]. Our co-culture of TERTp WT and MUT cells from the same tumor seems to recapitulate these traits better than monocultures, suggesting that interactions between TERTp MUT and WT cells may be at play.

Overall, our data show that STAR-FACS provides a reproducible platform for mutation-based cell labeling and separation. Our protocol generates high quality transcriptomic data that can be used to study phenotypic diversity driven by SNVs.

### TERTp mutation and single-cell transcriptomic heterogeneity

Recent advances in single cell transcriptomic profiling revealed the extent of intratumor heterogeneity in many cancer types, including GBM[34–38]. Genetic heterogeneity at the single cell level can be inferred from transcriptomic data and provide information about the copy number state of individual cells[39,40]. However, identifying point mutations in single-cell data remains challenging. Therefore, we tested whether STAR-FACS could be used to enrich cell population based on SNV to then perform a single-cell transcriptomic profiling of the mutant and wild-type populations.

Our fixation and STAR-FACS protocol was tested in combination with the split-pool single-cell barcoding protocol, commercialized by Parse Biosciences[41]. Evercode WT, based on SPLiT-seq[41], is a single-cell RNAseq method that labels the cellular origin of a transcript through combinatorial barcoding and is compatible with fixed cells and nuclei. Using an OS25-4 and OS23-2 co-culture (MIX A) and OS21-3 and OS21-8 co-culture (MIX B), we performed TERTp C228T STAR-FACS followed by the Evercode manufacturer’s protocol with several modifications (**Fig. 4a**, see **Methods** for details). Briefly, cells were processed through the STAR-FACS protocol as described above, sorted into TERTp WT and MUT fractions and frozen. Upon thawing, cells were counted, and each population was distributed into an Evercode well containing a well-specific barcode. At this step cDNA is generated using an *in cell* reverse transcription. Subsequently, the cells were pooled and distributed in 96 wells and an *in cell* ligation was performed to append a second well-specific barcode to the cDNA molecules. This step was repeated one more time to add a third-round barcode, which also includes a unique molecular identifier, followed by cell pooling, splitting into two sub-libraries, cell lysis, and cDNA extraction. Finally, the fourth sub-library-specific barcode was added in the last library preparation PCR. The advantage of the Evercode barcoding is that it allows for a broad range of input cell numbers, from a few hundred to over 100 million cells.

**Figure 4.**
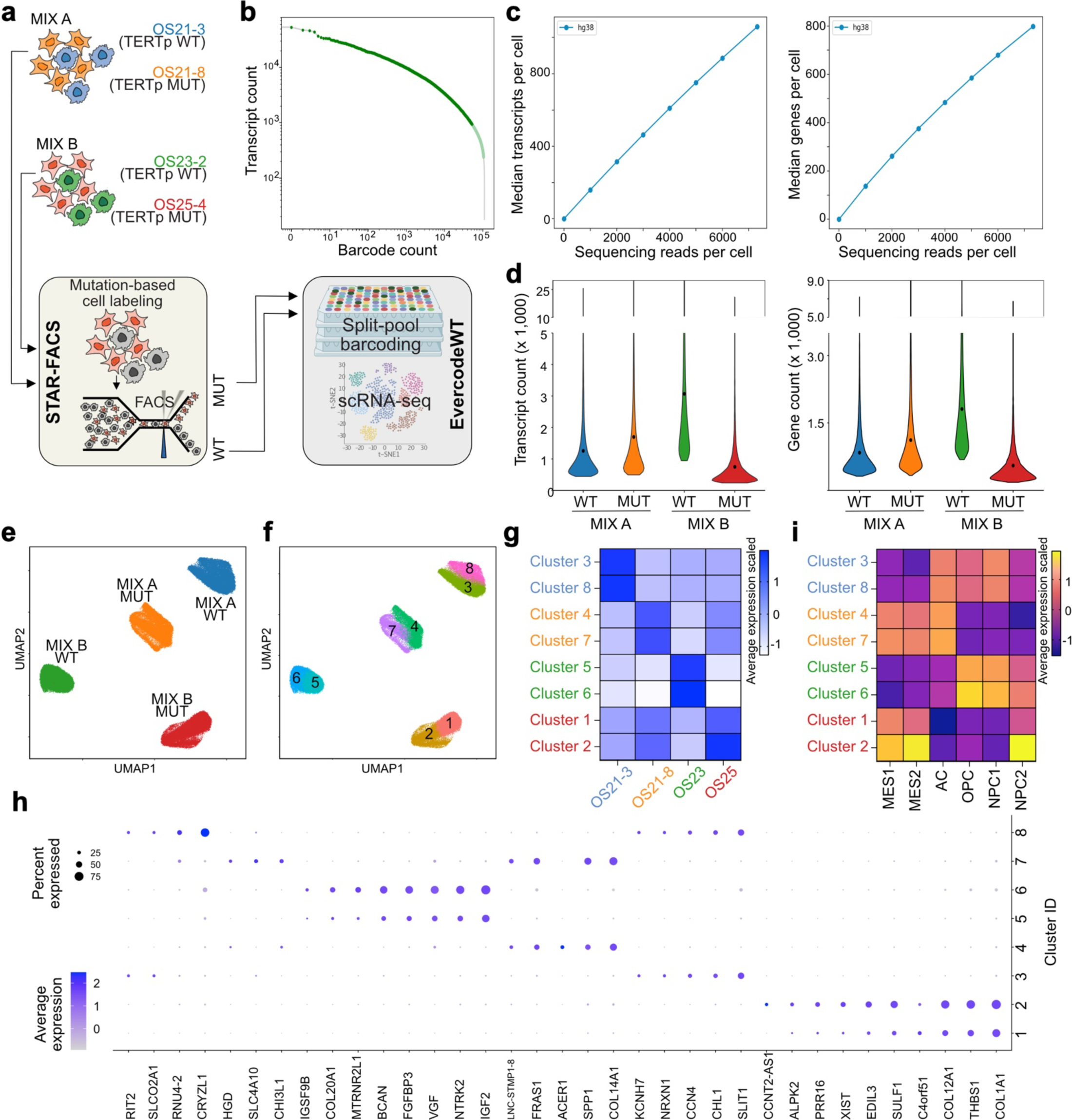
Single cell sequencing following TERT promoter mutation-based separation. **a**) Experimental outline. Two sets (MIX A and MIX B) of TERT promoter mutant (MUT) and wild-type (WT) cells were co-cultured for 10-14 days and separated by STAR-FACS. The fractions were subject to split-pool barcoding with Evercode protocol for single-cell RNA seq (scRNA-seq). **b**) Single-cell barcoding efficiency across all cells in combined MIX A and MIX B library. **c**) Single-cell sequencing depth for MIX A (left panel) and MIX B (right panel). The relatively low median transcript per cell is due to the high input cell number used in the barcoding step. **d**) Transcript count distribution in individual fractions sorted based on STAR-FACS from MIX A and MIX B co-cultures. **e**) Clustering of STAR-FACS-sorted single cells colored by the fraction of origin barcode. **f**) Clustering of STAR-FACS-sorted single cells colored by cluster. **g**) Average expression of parental line signature in single-cell clusters confirms the origin of the cells. **h**) Genes differentially expressed between the clusters. **i**) Average expression of GBM cell state-defining metamodules (Neftel et al., Cell 2019) in STAR-FACS-sorted clusters of cells. MES – mesenchymal-like program. AC – astrocyte-like program. OPC – oligodendrocyte-precursor-like program. NPC – neural-progenitor-like program.

We labeled 92,932 cells from our four subpopulations derived from MIX A and MIX B co-cultures (**Supplementary Table 1, Supplementary Fig. 4a, b**). High input cell number (10x over the manufacturer’s recommendation for the Mini kit) resulted in low sequencing saturation and relatively low transcript and gene detection per cell (**Fig. 4b-d**). However, the median count of 798 genes per cell is sufficient to perform clustering based on gene expression (**Fig. 4 d-g, Supplementary Fig. 4b**). UMAP dimensionality reduction clustered our cells into four major clusters, corresponding to the expected four populations (**Fig. 4 e, f**). The TERTp MUT subpopulations from MIX A and B are distinct from TERTp WT subpopulations and each one consists of two sub-clusters of cells, which are largely driven by changes in levels of cluster-specific markers. Bulk transcriptomic analysis confirmed the origin of the Evercode single-cell-based subpopulations (**Fig. 4g, Supplementary Fig. 4c**), proving that STAR-FACS can be successfully used in conjunction with single-cell transcriptomics. Interestingly, cluster 7 within the TERTp MUT cells from MIX A is distinct from cluster 4 (**Fig. 4h, Supplementary Fig. 4d**), expressing CHI3L1, HGD and SLC4A10, which are linked to stemness and neuronal excitability[42–44]. Moreover, analysis of gene signatures linked to single-cell transcriptional states in GBM revealed that both OS21-8 and OS21-3 derived fractions share levels of the astrocytic-like program (**Fig. 4i**; AC signature), but the TERTp mutant OS21-8 STAR-FACS-sorted cells are more mesenchymal (MES1 and MES2 signature). The mesenchymal state is also enriched in clusters 1 and 2 originating from OS25-4 TERTp mutant cells, but the cluster 2 subpopulation also expresses high levels of neural progenitor-like genes (NPC2 signature). These differences were not apparent from bulk sequencing, suggesting that even low-coverage single-cell RNAseq can help uncover traits unique to TERTp wild-type and mutant populations of cancer cells.

### Chromatin state profiling in TERTp mutation-sorted GBM cells

Since TERTp mutations affect the structure of the promoter and may have long-range effects on chromatin organization, we tested whether STAR-FACS TERTp mutant cell enrichment would allow for epigenetic profiling. Cleavage Under Targets & Tagmentation (CUT&Tag) is a recently developed approach that increases the signal-to-noise ratio of chromatin profiling by delivering chromatin marker-specific antibodies tethered to cut-and-paste transposase Tn5 into permeabilized cells or nuclei[45,46].

Activation of the transposase allows for simultaneous DNA cleavage and incorporation of adapters for paired-end DNA sequencing. We reasoned that our fixed and permeabilized, yet intact cells, after STAR-FACS could be used as input material for CUT&Tag. As a proof of principle, we performed CUT&Tag targeting histone H3 K27 acetylation (K27ac) and histone H3 K27 trimethylation (K27me^3^) in nuclei extracted from approximately 250,000 cells sorted for TERTp mutation with STAR-FACS from a mix of U87-MG and DK-MG cells. As a reference, we also performed CUT&Tag on fresh cells and nuclei from fixed cells. The chromatin landscapes for the STAR-FACS Alexa-647-sorted TERTp C228T mutant cells correspond to the fresh cells and fixed nuclei from the U87-MG GBM cell line, while the Cy3-sorted fraction of TERTp C250T mutant cells align with the fresh cells and fixed nuclei from DK-MG cells, confirming the identity of the sorted fractions (**Fig. 5a-c, Supplementary Fig. 5a**). The distribution of CUT&Tag signal at the transcription start sites is also similar between the post-STAR-FACS samples, fresh cell lines, and nuclei from fixed cell lines of origin (**Fig. 5b, Supplementary Fig. 5b**). Although the magnitude of the histone mark peaks in chromatin from fresh cell lines is higher compared to nuclei from fixed cells and STAR-FACS sorted samples, there is a strong overlap in chromatin regions captured in STAR-FACS, fixed nuclei and fresh cells reference (**Supplementary Fig. 5c-e, Supplementary Table 2**). Overall, these results confirm that mutation-based sorting is compatible with chromatin profiling by CUT&Tag.

**Fig. 5.**
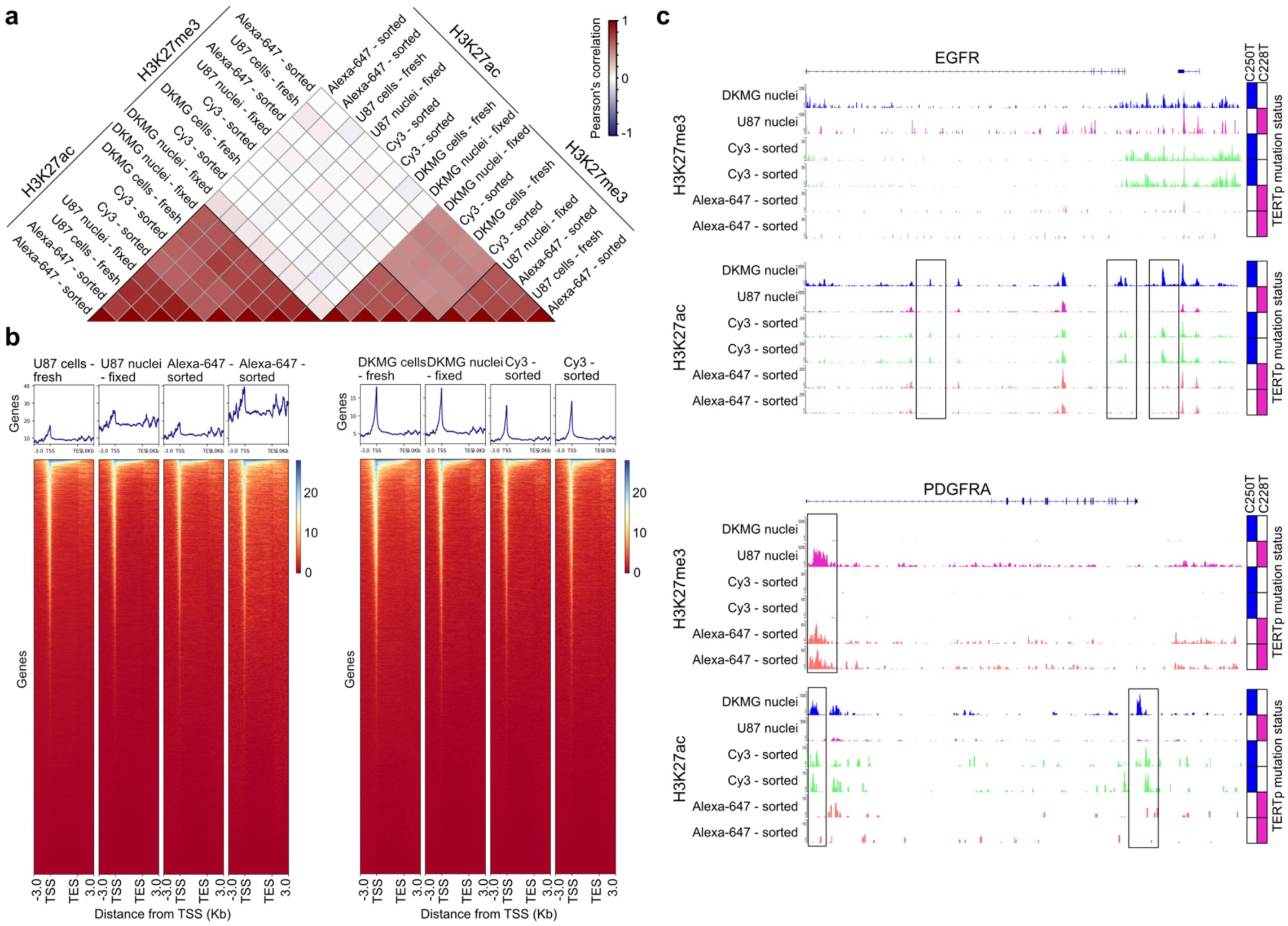
Chromatin state analysis with CUT&Tag after TERT promoter mutation-based cell separation. **a**) Correlation matrix for CUT&Tag results. Fresh U87-MG and DK-MG cells and nuclei from fixed cells and U87/DKMG mixed cells sorted by STAR-FACS were subject to CUT&Tag for H3K27me3 and H3K27ac histone marks. The genome was split into 500 bp bins, and a Pearson correlation of the log2-transformed values of read counts in each bin was calculated between the datasets. Alexa-647-sorted samples - C228Tmutant. Cy3-sorted samples – C250T mutant. **b**) H3K27ac mark distribution near transcription start sites (TSS) and transcription end sites (TES). **c**) Comparison of H3K27me3 and H3K27ac histone marks identified by CUT&Tag in two representative loci, EGFR and PDGFRA genes. Rectangles indicate areas of differential patterning for DK-MG and U87-MG cells. TERT promoter mutation status of the samples is shown on the sidebar: blue – C250T mutation, magenta – C228T mutation. Matching scaling was selected for control samples (DK-MG and U87-MG cells) and STAR-FACS sorted samples.

## DISCUSSION

Our study shows that STAR-FACS can be successfully used to separate cells based on a single nucleotide mutation status to perform transcriptomic and epigenetic profiling. We demonstrate that a model system of GBM primary neurosphere co-cultures containing a mix of TERTp MUT and WT cells can be effectively separated and profiled by bulk and single-cell transcriptomics. Despite *in cell* PCR and hybridization steps, STAR-FACS cells remain intact, and their transcriptomic profiles do not differ from those obtained from cells subject to standard 1% PFA fixation.

The *in cell* PCR step of our protocol can be performed to target two distinct point mutations at the same time. The two TERT promoter hotspot mutations, C228T and C250T are not commonly found together in the same tumor[30,47]. However, the ability to detect both hotspots simultaneously covers 80-90% of alterations in the promoter found in glioblastoma[30,48], increasing the chance of labeling a TERT mutant population without the necessity to know which specific hotspot is present. Dual mutation detection will also capture rare cases with C228T and C250T co-occurence[49]. Multiplexing STAR-FACS for the detection of several mutations at the same time might be challenging since the specificity of the assay depends on PCR. Achieving highly specific and efficient co-amplification of mutations located in genomic regions with vastly different GC content and secondary structure is inherently difficult and will limit the use of the method. Given that TERT promoter mutations are early events in GBM evolution, and that tumor purity can vary significantly between regions of the same tumor (from 20-90%)[48], STAR-FACS can enable enrichment in tumor purity for downstream applications. Furthermore, this method can also be used to assess subclonal populations. A recent report on TERT promoter mutation suggests that its subclonal frequencies are likely artifacts of tumor purity[48]. However, PCR bias towards the mutant allele of the TERT promoter, caused by decrease in GC content and possible unfolding of the G-quadruplex DNA structure[50], may contribute to Sanger sequencing and targeted sequencing underrepresentation of the wild-type allele. Since STAR-FACS is a single cell method, this bias would not impact the quantification of TERTp mutation frequency. As shown in our example of tumor OS21, non-clonal TERTp mutations can be detected and separated by STAR-FACS, providing new insight into evolutionary dynamics of highly heterogeneous tumors.

Our single-cell analysis of TERTp wild-type and mutant cells derived from the same tumor and co-cultured prior to the STAR-FACS-based separation indicates an increase in the mesenchymal transcriptional program in mutant cells. The epithelial-to-mesenchymal transition pathway is also overrepresented in bulk analysis of mutant cells compared to TERTp WT. Thus, TERTp mutations leading to reactivation of TERT expression in mutant cells provide two key benefits to the cells: overcoming telomere shortening and activating the pro-invasive mesenchymal phenotype. Interestingly, the TERTp WT cells derived from the same tumor have more neural-progenitor and oligodendrocyte-progenitor-like expression programs and high levels of anti-migratory SLIT1[51] and CCN4/Wnt-induced signaling protein 1, involved in maintaining glioma stem cells and supporting pro-tumor microenvironment[52]. These TERTp WT cells also upregulate the G2M checkpoint, which is linked to increased resistance to DNA damage by alkylating agents, such as temozolomide, the standard of care chemotherapeutic for GBM[53]. Therefore, distinct regions of tumor OS21 harbor cancer cells that may have diverged early in the tumor evolution and established opposing strategies to maintain the population, by either enhanced cellular motility or adopting progenitor-like phenotype. STAR-FACS will enable studies of such rare cell populations with genetically divergent traits in non-coding portions of the genome.

In our proof-of-concept experiments we demonstrate that STAR-FACS can be combined with the CUT&Tag assay for histone marks. The advantage of CUT&Tag is that it can be applied to a small number of cells (<5,000). Moreover, unlike ATAC-seq, the choice of antibodies for chromatin capture allows flexibility in studying transcription factors and chromatin remodeling proteins. Epigenetic landscapes are largely affected by the tissue environment. Thus, the interactions between the genetic and epigenetic heterogeneity in tumors may be lost or altered when studied *in vitro*. Methods such as spatial CUT&Tag[54], which allow for profiling of the chromatin state at cellular resolution in intact tissue, require custom-made microfluidics and are not easy to implement. Therefore, rapid tissue dissociation or nuclear extraction from formalin-fixed paraffin-embedded tissues are currently widely used to interrogate the epigenetic state of human tumor cells[55]. Future application of STAR-FACS to freshly dissociated human tumor tissue will enable us to decipher how point mutations alter global chromatin profiles, without the need to culture the cellular subpopulations of interest. STAR-FACS could also be used to enrich for cells with selected point mutations from CRISPR-edited population, providing an alternative to single-cell cloning.

## CONCLUSIONS

STAR-FACS is a novel cell sorting method allowing separation of cells based on a single nucleotide variation in their DNA. STAR-FACS-separated cells can be used in single-cell and bulk transcriptomic studies, as well as in chromatin state profiling by CUT&Tag. Thus, the application of STAR-FACS will enable connecting genetic, epigenetic and transcriptomic diversity in freshly dissociated tissues and heterogenous cell cultures, without the need of single cell cloning. The low cost and use of standard lab equipment in our STAR-FACS protocol makes it user-friendly and scalable.

## METHODS

### Reagents

All reagents used in the study are listed in **Supplementary Table 3**.

### Human tissue samples

All experiments with the use of human tumor tissue were approved by Scripps Research IRB protocol #IRB-18-7209 and Cleveland Clinic Martin Health surgical discards guidelines. GBM diagnosis was confirmed by fresh frozen histopathology at the hospital. Fresh GBM tumor tissue was collected directly from the operating room at the time of surgery. For each tumor sampling, MRI navigation (Styker) was used to guide the collection, enabling us to spatially localize the resections. Tumor fragments were dissociated mechanically, saving a piece for FFPE embedding, and enzymatically with papain-based Brain Tumor Dissociation Kit P (Milteny Biotec). Cell lines were generated from ∼1 x 10^6^ enzymatically dissociated cells or from a ∼5mm^3^ piece of mechanically dissociated tissue, as previously described[56,57]. Red blood cells were removed from enzymatically dissociated cells by cell pellet incubation in RBC lysis buffer (155 mM NH_4_Cl, 12mM NaHCO_3_, 0.1 mM EDTA) for 5 minutes at room temperature. Cell cultures derived by mechanical dissociation of the tissue, as opposed to enzymatic dissociation described above, were treated with RBC lysis within 3 days from starting the culture.

OS11-5 neurosphere culture was derived from case OS11 GBM (60 year old, female); OS21-3 and OS21-8 cultures were derived from case OS21 GBM (60 year old, male); OS23-2, OS23-3, OS23-4, OS23-5 were derived from case OS23 GBM (74 year old, male); OS25-4 was derived from case OS25 GBM (75 year old, female).

### Cell cultures

U-87 MG cell line (ATCC) was cultured in DMEM (Corning) with 10% FBS (Avantor) according to manufacturer’s protocol. DK-MG cell line (DSMZ) was cultured in RPMI with 2mM L-glutamine (HyClone) with 10% FBS (Avantor) according to the manufacturer’s protocol.

Primary GBM cell lines were cultured at 1 million cells per flask in T25 flasks with 5ml of Neurobasal media (Life Technologies), with 0.5 % B27 Supplement (Life Technologies), 0.5% N2 Supplement (Life Technologies), 1% GlutaMAX Supplement (FisherScientific), 20ng/ml EGF (Milteny Biotec), 20ng/ml FGF (Arcobiosystems) and 1% PenStrep (Fisher Scientific). Growth factors were added every 2 days. Accutase (Fisher Scientific) was used to dissociate the spheres and passage the adherent cells. All primary cell lines were used between passage 6 and 10.

All cell lines were maintained in humidified incubators at 37°C, 4% CO_2_. Cells were routinely checked for mycoplasma contamination using Mycoplasma PCR Detection kit (Applied Biological Materials).

For co-culture experiments, equal cell numbers from two GBM cell lines were seeded together and cultured for 10-14 days.

TERT promoter mutation status of each cell line was confirmed by Sanger sequencing.

## STAR-FACS - mutation-based cell sorting

### Brief overview

Fixed and permeabilized intact cells are subject to two rounds of PCR (see below for details). During the first round of *in cell* PCR, a 597bp amplicon was generated to amplify the TERT promoter region of interest and to provide an abundant template for the allele-specific barcoding performed with the 2nd round of *in cell* PCR. In the 2nd round of in cell PCR allele-specific primers are used to specifically amplify the C228T or the C250T mutant allele and to add a unique barcode to these amplicons. Next, a hybridization step with a fluorescent probe complementary to the unique barcode is performed to fluorescently label cells harboring a specific allele.

The primers used in both the 1st and 2nd rounds of *in cell* PCR were biotinylated to enable the removal of excess amplicon that may interfere with downstream profiling of the labeled cells.

### Fixation

The protocol detailed here is suitable for up to 1 x 10^6^ total cells and it has been validated with as few as ∼3 x 10^5^ cells of sizes ranging from 9 µm to 26-28 µm. For each of the samples processed, 3-5 x 10^6^ cells were used as initial input and ∼80% of cells were recovered after *in cell* labeling and served as input for cell sorting.

After harvesting and mixing the U-87 MG and DK-MG cells, they were centrifuged at 4°C, 200g for 5 minutes, while the primary OS cells were centrifuged at 4°C, 200g for 7 minutes. The cell pellet was resuspended in 2 mL of cold PBS-RI-PI (1x PBS + 0.05U/μL SUPERase•In RNase Inhibitor (ThermoFisher Scientific) + 1 tablet of Complete protease inhibitor cocktail/50mL (FisherScientific)) and kept on ice. The cells were filtered through a 40 μm strainer into a 50 mL falcon tube. 6 mL of ice cold 1.33% methanol and RNase-free paraformaldehyde solution (PFA in 1x PBS-RI-PI) were then added to the 2 mL single cell suspension so that the final concentration of paraformaldehyde in the mixture equaled 1%. The tube was gently inverted three times and placed on ice for 10 minutes. Then, 160 μL of 10% Triton X-100 was added and the tube was gently inverted three times before being placed on ice for additional 3 minutes. Crosslinking was stopped by adding 8 mL of 1.25M Glycine (in RNase, DNase and Proteinase free water) to the mixture and the tube was gently inverted three times. The fixed and permeabilized single cell suspension was centrifuged at 4°C, 400g for 3-5 minutes and the supernatant was aspirated. The cells were washed with 1 ml of ice-cold PBS-RI-PI and pelleted at 4°C, 400g for 3-5 minutes.

Subsequently, the cell pellet was resuspended in 1.9 mL of ice-cold PBS-RI-PI and filtered through a 40 µM strainer until not large clumps were visible. 33 µl of DMSO were added to the cell suspension and the tube was flickered to mix and incubated on ice for 1 minute. Additional 33 µl of DMSO was added followed by flickering and 1 minute incubation on ice. Finally, 34 µl of DMSO was added, and the single cell solution was mixed by gentle pipetting before being aliquoted in 2 cryovials (not to exceed 5 x 10^6^ cells/vial) and cryopreserved at −80°C overnight.

A variant of this protocol was employed to test the compatibility of STAR-FACS with chromatin profiling via CUT&TAG. To avoid epitope masking due to excessive crosslinking, the concentration of paraformaldehyde solution added to the sample was reduced to 0.33% so that the final concentration of paraformaldehyde in the cell suspension equaled 0.250%. The Glycine neutralization step was scaled accordingly.

### Primer design for allele-specific in cell PCR

All primers (**Supplementary Table 4**) were designed based on the GRCh38 genome assembly, release 104. Furthermore, all the primers discussed here were biotinylated at the 5’ end to enable the removal of the amplicon from the labeled cells before downstream profiling. The first round of *in cell* PCR was intended to amplify a small region of the TERT promoter around the two hotspot mutations, C228T and C250T, targeted for the allele-specific cell labeling. For this round of *in cell* PCR, and an array of 4 forward primers and 7 reverse primers were generated with primer3[58], all the combinations were tested using 100ng and 900ng of DNA extracted from the same cells used in the labeling experiment. The amplicons generated were gel extracted with Qiagen QIAquick Gel Extraction Kit according to the manufacturer’s recommendations and profiled by Sanger sequencing to test specificity.

For the primer combination selected (TERT_F1: TTCGACCTCTCTCCGCTGG, TERT_R1: CAGGGCACGCACACCAG) we used a reaction mix with 1x modified OneTaq GC Reaction Buffer (80 mM Tris-SO_4_, 20 mM (NH_4_)_2_SO_4_,1.5 mM MgCl_2_, 0.06% IGEPAL CA-630, 0.05% Tween 20, 5% Glycerol, pH 9 at 25°C), 400 mM Betaine, 200µM dNTPs, 0.2µM TERT_F1 primer, 0.2µM TERT_R1 primer, 0.75 U/30µL reaction of OneTaq Hot Start DNA polymerase. After initial incubation at 94°C for 2 min, the reaction was cycled 5-10 times at 94°C for 20 sec, 56°C for 20 sec, 68°C for 40 sec, followed by a final elongation step at 68°C for 5 min and hold at 4°C. The amplicon generated was 597bp long and the approach was further validated using DNA extracted from fixed cells as input to make sure that DNA fragmentation due to the fixation did not interfere with the amplification reaction.

For the second round of in cell PCR, first two allele specific forward primers were designed and one universal reverse primer (TERT_C228T_F: TGTCGACGCAAAACCGGTTCTCCCCGGCCCAGCCCTTT, TERT_C250T_F:TCACCAGGCAATAGCCGTTCCCCCGTCCCGACCCCTTG, TERTall_R2_univ:GCGATATGACGACGCGAATAGGCTTCCCACGTGCGCAGC.) For the C250T mutation, the mutant allele was the second nucleotide from the 3’ end, and an additional mismatch was added at the first nucleotide from the 3’. For the C228T mutation the mutant allele was the second nucleotide from the 3’ end, and an additional mismatch was added at the third nucleotide from the 3’ end. The mismatch helped improve the specific amplification of the targeted allele.

Three barcodes foreign to the human genome were designed using unwords[59] and manually edited to make them compatible with the TERT_C228T_F, TERT_C250T_F and TERTall_R2_univ primers. NUPAK[60] was used to verify that the full primers did not form any secondary structure that would hinder their ability to amplify the desired targets. The same PCR conditions were optimized to be used with the TERT_C228T_F and the TERT_C250T_F primer. The reaction mix was prepared with 1X modified OneTaq GC Reaction Buffer (80 mM Tris-SO_4_, 20 mM (NH_4_)_2_SO_4_,1.5 mM MgCl_2_, 0.06% IGEPAL CA-630, 0.05% Tween 20, 5% Glycerol, pH 9 at 25°C), 200µM dNTPs, 0.2µM TERT_C228T_F or TERT_C250T_F, 0.2µM TERTall_R2_univ primer, 0.75 U/30µL reaction of OneTaq Hot Start DNA polymerase. After initial incubation at 94°C for 2 min, the reaction was cycled 15 times at 94°C for 20 sec, 63°C for 20 sec, 68°C for 20 sec, followed by a final elongation step at 68°C for 5 min and holding at 4°C. The TERT C250T PCR produced an amplicon of 181bp and the TERT C228T PCR produced an amplicon of 159 bp. Both were validated using DNA extracted from fixed cells with known TERT mutations and confirmed by Sanger sequencing.

The fluorescent probes were designed to match the unique barcode added to each mutation-specific forward primer. The fluorophore (Alexa-647 for C228T and Cy3 for C250T) was added at the 5’ end of the probe. Furthermore, the 3 nucleotides at the 5’ end and the 3 nucleotides at the 3’ end of the probes were replaced with LNAs in order to increase their rigidity and create a stronger bond with the targets.

### In cell specific-to-allele PCR

The fixed cells stored at −80°C were thawed at 37°C and once the ice crystals were almost completely dissolved, 1 mL of PBS-RI-PI was added at room temperature and the cells were centrifuged at 400g for 5 minutes. The cell pellet was washed once with an additional 1 ml of PBS-RI-PI and the cells were counted with an automatic cell counter or a hemocytometer. If large clumps were observed, the cells were passed through a 40 µm strainer. After centrifugation at room temperature the cells were re-suspended in the 1st round of *in cell* PCR master mix, 1X modified OneTaq GC Reaction Buffer (80 mM Tris-SO_4_, 20 mM (NH_4_)_2_SO_4_,1.5 mM MgCl_2_, 0.06% IGEPAL CA-630, 0.05% Tween 20, 5% Glycerol, pH 9 at 25°C), 400 mM Betaine, 200µM dNTPs, 0.2µM TERT_F1 primer, 0.2µM TERT_R1 primer, 0.75 U/30µL reaction of OneTaq Hot Start DNA polymerase. A total of 32 (30µL) reactions were prepared for each cell mixture processed, and each reaction contained ∼150,000 fixed cells.

After initial incubation at 94°C for 2 min, the reactions were cycled 10 times at 94°C for 20 sec, 56°C for 20 sec, 68°C for 40 sec, followed by a final elongation step at 68°C for 5 min. The next step on the thermocycler was set to hold the samples at 30°C. As soon as the samples reached the holding temperature, they were removed from the thermocycle and the reactions were pooled and centrifuged at room temperature, 400g for 5 minutes. The supernatant was removed, and the cell pellet was resuspended in the 2nd round of *in cell* PCR master mix, 1X modified OneTaq GC Reaction Buffer (80 mM Tris-SO_4_, 20 mM (NH_4_)_2_SO_4_,1.5 mM MgCl_2_, 0.06% IGEPAL CA-630, 0.05% Tween 20, 5% Glycerol, pH 9 at 25°C), 200µM dNTPs, 0.2µM TERT_C228T_F (for the simultaneous labelling of the C228T and C250T mutation an equal mix of 0.2 µM TERT_C228T_F and TERT_C250T_F was added), 0.2µM TERTall_R2_univ primer, 0.75 U/30µL reaction of OneTaq Hot Start DNA polymerase. The mixture was split in a total of 32 (30 µL) reactions and placed in the thermocycler holding at 30°C. The holding step was manually skipped and after the initial incubation at 94°C for 2 min, the reactions were cycled 15 times at 94°C for 20 sec, 63°C for 20 sec, 68°C for 20 sec, followed by a final elongation step at 68°C for 5 min and holding at 4°C.

### Fluorescent probe hybridization

The 32 PCR reactions from the 2nd round of in cell PCR were pooled and centrifuged at room temperature for 5 minutes, 400g. The supernatant was removed, and the cells were washed once with PBS-RI-PI. Then, the cell pellet was dehydrated through three washes with 75% EtOH, 80% EtOH and 100% EtOH. In between each wash, the pellet was spun down at room temperature 400g for 3 minutes. The cell pellet was dried at room temperature for 10-15 minutes.

The hybridization mix (sufficient for a total of ∼1 x 10^6^ cells) was prepared with 215.31 µL of hybridization buffer (2x SSC, 20% formamide, 10% dextran sulfate), 25.9 µL of 1mg/mL human COT1 DNA (Life Technologies), 2.59 µL of 10 mg/mL shredded salmon sperm DNA (Life Technologies; both COT1 and salmon sperm DNA were biotinylated with EZ-Link Psoralen-PEG3-Biotin via UV activated intercalation prior to their use in the buffer to facilitate their removal when preparing the labelled cells for sequencing) and 6.2 µL mutation specific fluorescent probe (the for the simultaneous labeling of the C228T and C250T mutation, 4 µL of each probe were used). The hybridization mix was further supplemented with 0.05U/µL of SUPERase•In RNase Inhibitor (ThermoFisher).

The dried cell pellet was resuspended in the hybridization mix prewarmed at 37°C and incubated at 74°C for 7 minutes followed by overnight incubation (10-12 hours and never to exceed 15 hours) at 37°C, protected from light.

Next, 1 mL of 0.4x SSC, 0.3% NP-40 (in 1X PBS-RI-PI) was added to the cells in hybridization buffer, mixed by gentle pipetting up and down three times and centrifuged at room temperature, 600g for 5 minutes. The supernatant was removed, and the pellet was resuspended in 1 mL 0.4x SSC, 0.3% NP-40 (in 1X PBS-RI-PI) preheated at 74°C. The cells were resuspended by gentle pipetting up and down 3 times and centrifuged at room temperature, 400g for 3 minutes. The supernatant was removed, and the cells were resuspend in 2x SSC, 0.1% NP-40 (in 1X PBS-RI-PI) and centrifuged at room temperature, 400g for 3 minutes. A fourth wash was performed with 2x SSC (in 1X PBS-RI-PI) and the cells were centrifuged at room temperature, 400g for 3 minutes. A final wash was performed with 1mL of 1X PBS-RI-PI, and the cells were resuspended in 1X PBS-RI-PI, 0.5mM EDTA and filtered through a 35 µm strainer into a flow cytometry tube.

### Cell sorting

Cells were sorted on BD FACSAria Fusion in 1X PBS-RI-PI, 0.5mM EDTA. Gating was set based on a positive control, U87 human glioblastoma cells, *TERT* promoter C228T mutation, or DK-MG, *TERT* promoter C250T mutation. Cell sorting was performed until ∼3 x 10^5^ cells were collected for each targeted fraction. The sorted cells were frozen in 1X PBS-RI-PI, 5% DMSO, and stored at −80°C for ∼2 days before proceeding with the downstream application.

### Bulk RNA extraction, sequencing and analysis

For bulk RNA-Seq, RNA was extracted from cell line pellets with TRIzol Reagent (ThermoFisher) and Direct-zol RNA Mini-Prep Plus kit (Zymo Research). Cells subject to STAR-FACS were thawed at 37°C and once the ice crystals were almost completely dissolved, 1 mL of PBS-RI-PI was added at room temperature and the cells were centrifuged at 400g for 5 minutes. The cell pellet was then resuspended in 200 µL of PBS-RI, then 200 µL of 2X lysis buffer (20 mM Tris pH 8.0, 400mM NaCl, 100 mM EDTA pH 8.0 and 4.4% SDS) was added and 40 µL of 20 mg/mL proteinase K (Life Technologies). The cells mixture was then incubated at 55°C for 2 hours with shaking at 400rpm.

Afterwards, the 400 µL cell lysate was diluted with 4 mL of Trizol and mixed by vortexing. The 5 mL of 100% EtOH was added to the sample in Trizol and again mixed by vortexing. Then the mixture was incubated at room temperature for 10 minutes on a nutator. The mixture was split loaded on two Zymo-Spin IICR columns and RNA purification with on-column DNase I treatment was carried out according to manufacturer’s recommendation. The RNA was eluted in RNase free water and the elution from both columns was combined.

The residual amplicon generated during the *in cell* PCRs and residual COT1 DNA and salmon sperm DNA leftover from the hybridization step were removed via Streptavidin pulldown from the purified RNA samples. Dynabeads MyOne Streptavidin C1 beads (Life Technologies) were washed for RNA application according to the manufacturer’s recommendation. The beads were then added to the samples and incubated at room temperature for 10-15 minutes on a nutator to allow the binding of the biotinylated DNA on the streptavidin beads. The biotinylated DNA was then separated with a magnet for ∼2 minutes and the unbound fraction with the RNA was reprecipitated with a standard EtOH precipitation with a new Zymo-Spin IICR column.

Libraries were prepared using NEBNext Ultra 2 directional kit with rRNA depletion module 1 (New England Biolabs) and sequenced according to the standard protocol. Paired-end 50×50 bp sequencing was performed on the Illumina NextSeq 2000 platform, yielding ∼20 million reads per library. Reads were aligned to the Homo sapiens GRCh38 release 104 reference genome with STAR aligner, version 2.7.9a[61]. Raw counts were computed with HTSeq, version 2.0.3[62].

Pseudo-alignment and transcript quantification was performed using Salmon[63]. Differential gene analysis was performed using DESeq2 R package[64].

To test whether remaining STAR-FACS amplicon was contributing to RNAseq data, the reads were aligned to a reference generated from the sequence corresponding to the amplicon. No significant overlap was found.

### Single-cell RNA sequencing and data analysis

Single-cell RNA seq was performed on two primary neurosphere co-cultures: MIX A, containing OS21-3 (TERTp WT) and OS21-8 (TERTp MUT) cells, derived from the same tumor; MIX B, containing OS23-2 (TERTp WT) and OS25-4 (TERTp MUT) cells. Cell and library preparation for scRNA-seq was performed according to Evercode Whole Transcriptome Mini protocol (Parse Biosciences), with a modification to exclude STAR-FACS amplicon from library preparation. To achieve this, a pool of six oligonucleotides with 3’ C3 spacer (forward and reverse), spanning the amplicons produced through the STAR-FACS labeling, was added to each sample prior to aliquoting it into the *in situ* reverse transcription plate (final concentration of oligos was 1.4uM). After that standard protocol was applied.

The reverse transcription plate was loaded as follows: wells 1, 2 and 3 - Alexa-647 positive fraction (TERTp MUT) of MIX A; wells 4, 5 and 6 - Alexa-647 negative fraction (TERTp WT) of MIX A; wells 7, 8 and 9 - Alexa-647 positive fraction (TERTp MUT) of MIX B; wells 10, 11 and 12 - Alexa-647 negative fraction (TERTp WT) of MIX B. The scRNA-Seq was targeting 2,500 cells per sample.

Demultiplexing and alignment of the data were performed using Parse Biosciences pipeline version 1.2.1 (available online from Parse Biosciences), the reads were aligned to *Homo sapiens* GRCh38 release 104 reference genome. The output of the pipeline was imported into R 4.3.3 with Seurat package 5.0.3[65], using ReadParseBio() function. The data was subset to include cells with 200-2500 transcripts and <5% mitochondrial gene content and normalized using “LogNormalize” method. Louvain clustering was performed using FindNeighbors() and FindClusters() functions with a resolution at 0.5.

### CUT&Tag

Cut&Tag was performed with CUT&Tag-IT Assay Kit, Anti-Rabbit (Active Motif) for the histone H3K27me3 and H3K27ac. For each CUT&Tag reaction 250,000 cells were used as input. One set of controls was prepared with freshly harvested U87-MG and DK-MG cells, processed according to the manufacturer’s recommendation with overnight primary antibody incubation at 4°C. A second set of controls was prepared with nuclei extracted from 0.25% PFA-fixed U87-MG and DK-MG cells, processed with the STAR-FACS protocol described above up to the *in cell* PCR step. Nuclei extracted from 1,000,000 cells resuspended in ½ volume ice-cold NE1 buffer (1 ml 1M HEPES-KOH pH 7.9, 500 μL 1M KCl, 12.5 μL 2 M spermidine, 500 µL 10% Triton X-100, and 10 ml glycerol in 38 ml nuclease-free H2O, supplemented with 1 Roche cOmplete protease inhibitor cocktail EDTA-Free) with light vortexing and incubated for 10 minutes on a nutator in the cold room. After centrifugation at 4°C, 1000 xg for 5 minutes, the nuclei were resuspended in CUT&Tag-IT wash buffer. 200-250,000 nuclei were aliquoted and used as input for the CUT&Tag reactions. The Active Motif protocol was performed according to the manufacturer’s recommendation, except for digitonin, which was omitted from the buffer preparation, as PFA fixed cells and nuclei were sufficiently permeabilized to allow antibody delivery. The primary antibody incubation was performed overnight at 4°C.

The experimental samples derived from the STAR-FACS labeling and sorting of U87-MG and DK-MG mixes (Alexa-647-sorted and Cy3-sorted, n=2 each) were processed using the same approach as the second set of controls. The nuclei were extracted from the sorted cells and the CUT&Tag protocol was performed with the modifications described above. 200-250,000 nuclei were targeted.

All sequencing libraries were prepared according to the manufacturer’s recommendations. Paired-end 50×50 bp sequencing was performed on the Illumina NextSeq 2000 platform, yielding ∼20 million reads per library. The data were first trimmed to remove adapters using Trim Galore version 0.6.5 (doi: 10.5281/zenodo.7598955) and standard parameters. Trimmed reads aligned to the Homo sapiens GRCh38 release 104 reference genome with Bowtie2 version 2.4.5[66] and the following options, --local --very-sensitive --no-mixed --no-discordant --phred33 -I 10 -X 700 -x. Then the reads were aligned to the Ecoli genome strain K12, substrain MG1655, with the following options, --end-to-end --very-sensitive --no-overlap --no-dovetail --no-mixed --no-discordant --phred33 -I 10 -X 700 -x. Properly paired, quality-filtered reads were extracted, indexed, and sorted with samtools version 1.19.2[67] and Picard version 3.1.1 (https://broadinstitute.github.io/picard/). Counts were scaled according to the scaling factor S = 10000/number of reads aligning to the E. coli genome. These reads correspond to the bacterial DNA carried over from pA-Tn5 purification and are used as a calibration standard instead of a spike-in for comaring samples in an experiment[45]. Tracks were generated with bedtools version 2.30.0[68] genomeconv command. Mitochondrial reads were removed, and peaks were called with MACS2 version 2.2.7.1[69] and the following options, macs2 callpeak -t <fragments.bed> -f BEDPE -g hs --keep-dup all -p 1e-5 -n <PEAKS> --SPMR.

Heatmaps for chromatin features over transcriptional units and CUT&TAG peaks were generated with deeptools version 3.5.2[70].

## DECLARATIONS

### Ethics approval and consent to participate

All experiments with the use of human tumor tissue were approved by Scripps Research IRB protocol #IRB-18-7209 and Cleveland Clinic Martin Health surgical discards guidelines.

### Consent for publication

Not applicable.

### Availability of data and materials

All transcriptomic and CUT&TAG sequencing data from this manuscript are available at NCBI SRA (accession number PRJNA1100531).

### Competing interests

The authors declare no competing financial interests. M.J. is a member of a scientific advisory board at ResistanceBio.

## Funding

This work was supported by the NIH K99/R00 CA201606 (M.J.), Florida Center for Brain Tumor Research Seed Grant (M.J.) and start-up funds from the Scripps Research Institute (M.J.).

## Authors’ contributions

R.S. and M.J conceived the study. R.S. performed all the experimental work and data analysis. Computational analysis was performed by R.S with M.F. Flow cytometry data analysis was performed by R.S with A.R.T. Y.W. helped with flow cytometry experiments. S.T. helped with single-cell sample preparation. O.S. collected the human surgical specimen and annotated clinical data. M.J. supervised the study. All authors contributed to writing the manuscript.

## Acknowledgments

We thank the members of the Janiszewska lab for their critical reading of this manuscript and useful discussion. We thank the staff at The Wertheim UF Scripps Institute Genomics Core, Flow Core, Bioinformatics and Statistics Core, and University of Florida ICBR NextGen Sequencing Core for their expert technical support. We thank Dr. Steven Henikoff for his advice on adapting CUT&Tag to STAR-FACS workflow.

