## Supplemental Figures for "Cell sorting based on single nucleotide variation enables characterization of mutation-dependent transcriptome and chromatin states"

### SUPPLEMENTARY FIGURES

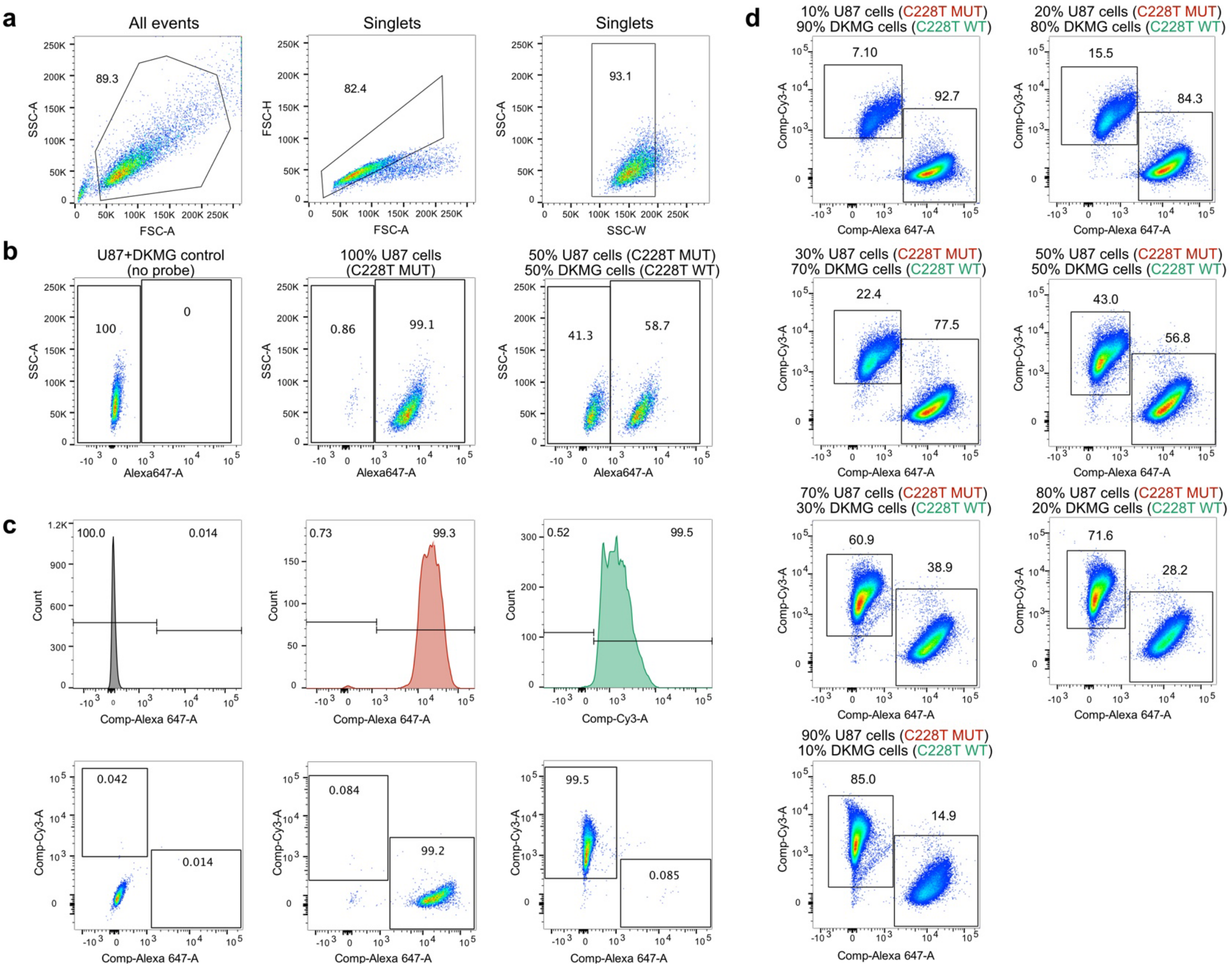


**Supplementary Figure 1.** Gating strategy, compensation controls and quantitative validation of STAR-FACS. **a**) FACS gating example for STAR-FACS labeled sample. **b**) STAR-FACS labeling for C228T mutation with Alexa-647 probe. Left panel – no probe control with 1:1 mix of U87-MG and DK-MG cells. Middle panel – 100% U87-MG cells. Right panel – 1:1 mix of U87-MG and DK-MG cells. **c**) Compensation controls for STAR-FACS labeling two TERTp hotspot mutations. C228T mutation-specific probe is labeled with Alexa-647 and C250T probe is labeled with Cy3. Upper panels are single color histograms of the data presented in lower panels. Left panels – no probe control. Middle panels– U87-MG cells (C288T mutant). Right panels - DK-MG cells (C250T mutant). **d**) STAR-FACS labeling two TERTp hotspot mutations on U87-MG and DK-MG cell mixtures. C228T mutation-specific probe is labeled with Alexa-647 and C250T probe is labeled with Cy3. Percentage of cells in corresponding gates is shown.

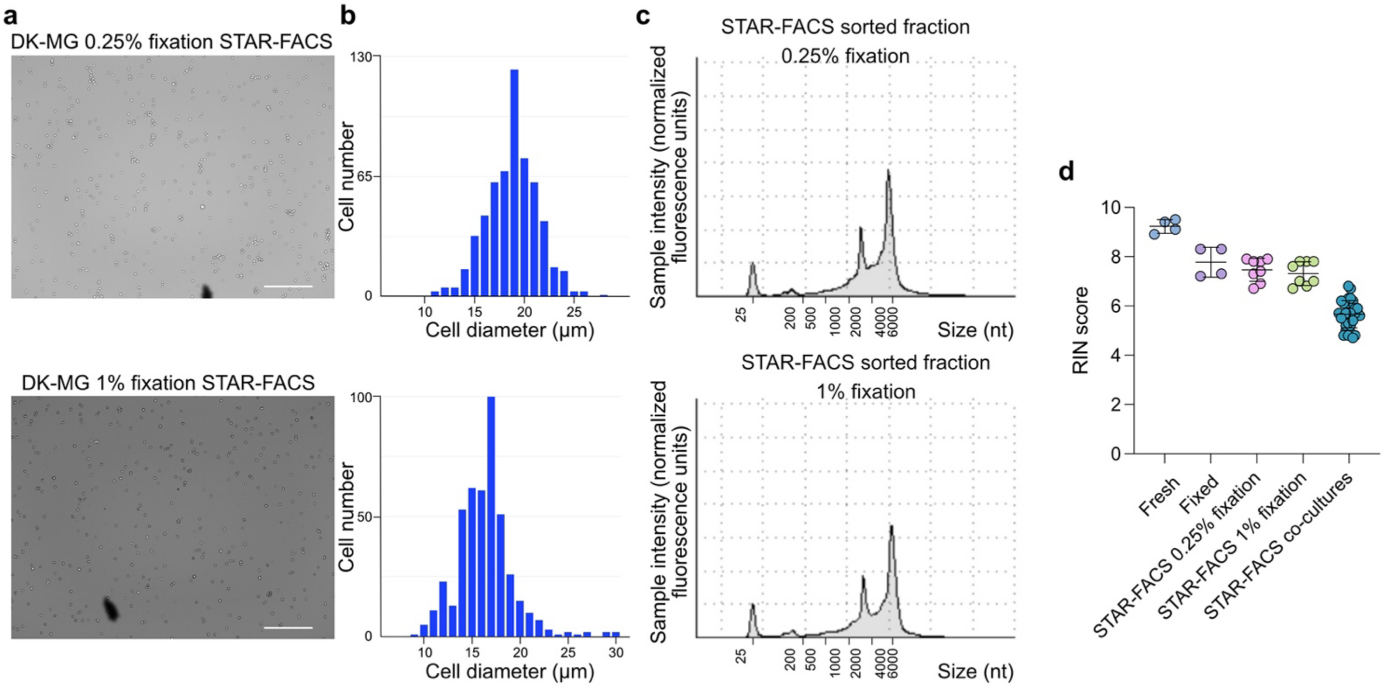


**Supplementary Figure 2.** Cell integrity and RNA quality post-STAR-FACS. **a**) Representative images of cell integrity after STAR-FACS with 0.25% (upper panel) and 1% fixation (lower panel). Scale bars - 500um. **b**) Cell diameter measurement corresponding to images in a). **c**) Representative TapeStation RNA size analysis. RNA extracted from STAR-FACS sorted fractions with 0.25% or 1% fixation protocol is shown. **d**) RNA integrity (RIN) scores of RNA extracted from freshly collected cells (n=4), 1% PFA fixed cells (n=4) and STAR-FACS-sorted cells (n=32).

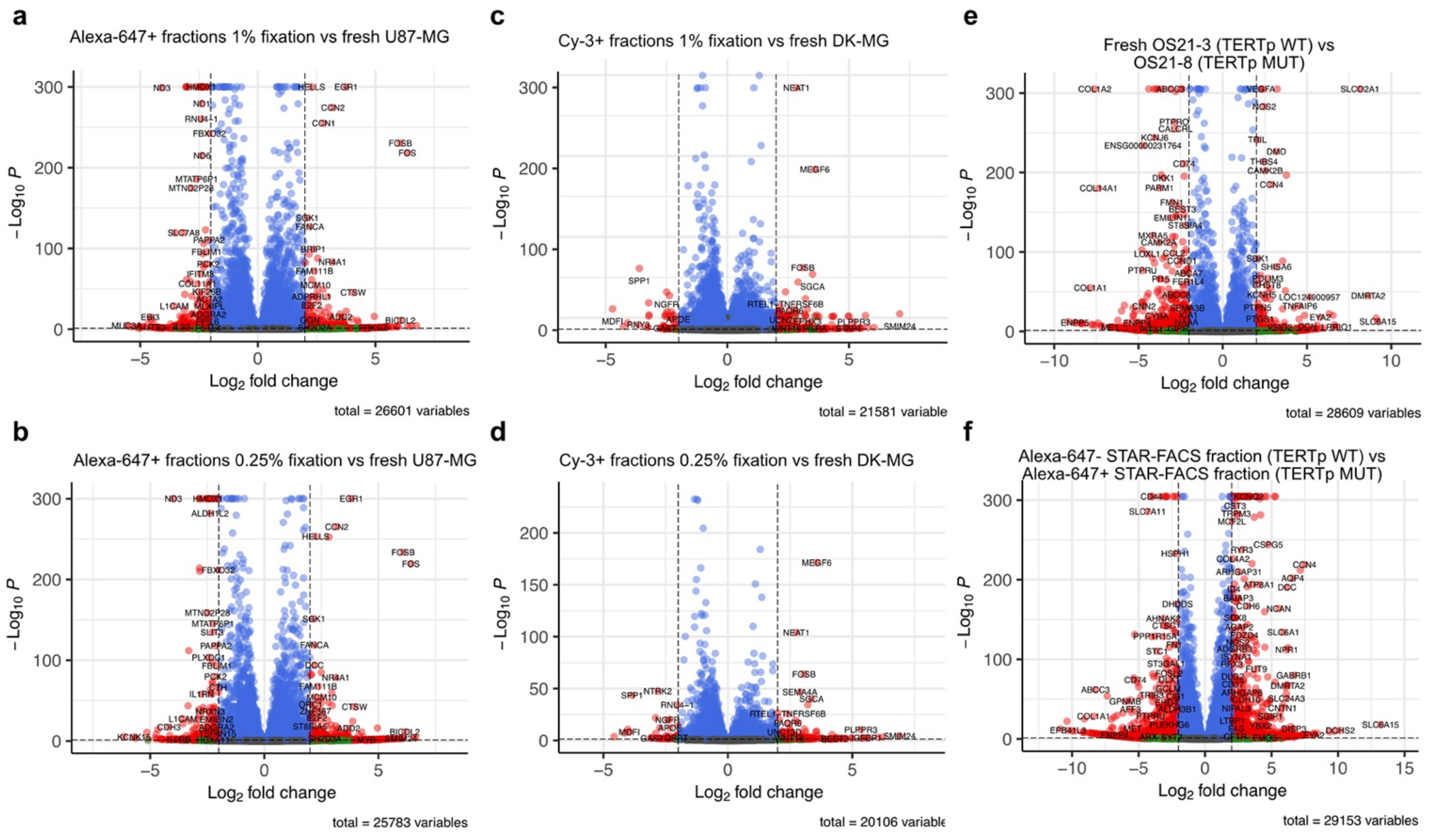


**Supplementary Figure 3.** Effect of fixation and STAR-FACS on differential gene expression analysis. **a-d**) Comparison of gene expression in STAR-FACS sorted cell fractions with parental cell line. U87-MG (TERTp MUT) cell were mixed with DK-MG cells (TERTp WT) and STAR-FACS sorted. Alexa-647+ correspond to U87-MG cells and Cy-3+ fractions correspond to DK-MG cells. Fixation of 1% was performed in **a**) and **c**) while **b**) and **d**) show results for 0.25% fixation. **e**) Differential gene expression between OS21-3 and OS21-8 freshly collected neurospheres. **f**) Differential gene expression between STAR-FACS TERTp positive and negative fractions from OS21-3 and OS21-8 co-culture experiment. OS21-3 (TERTp WT) corresponds to Alexa-647- fraction, while OS21-8 (TERTp MUT) corresponds to Alexa-647+ fraction.

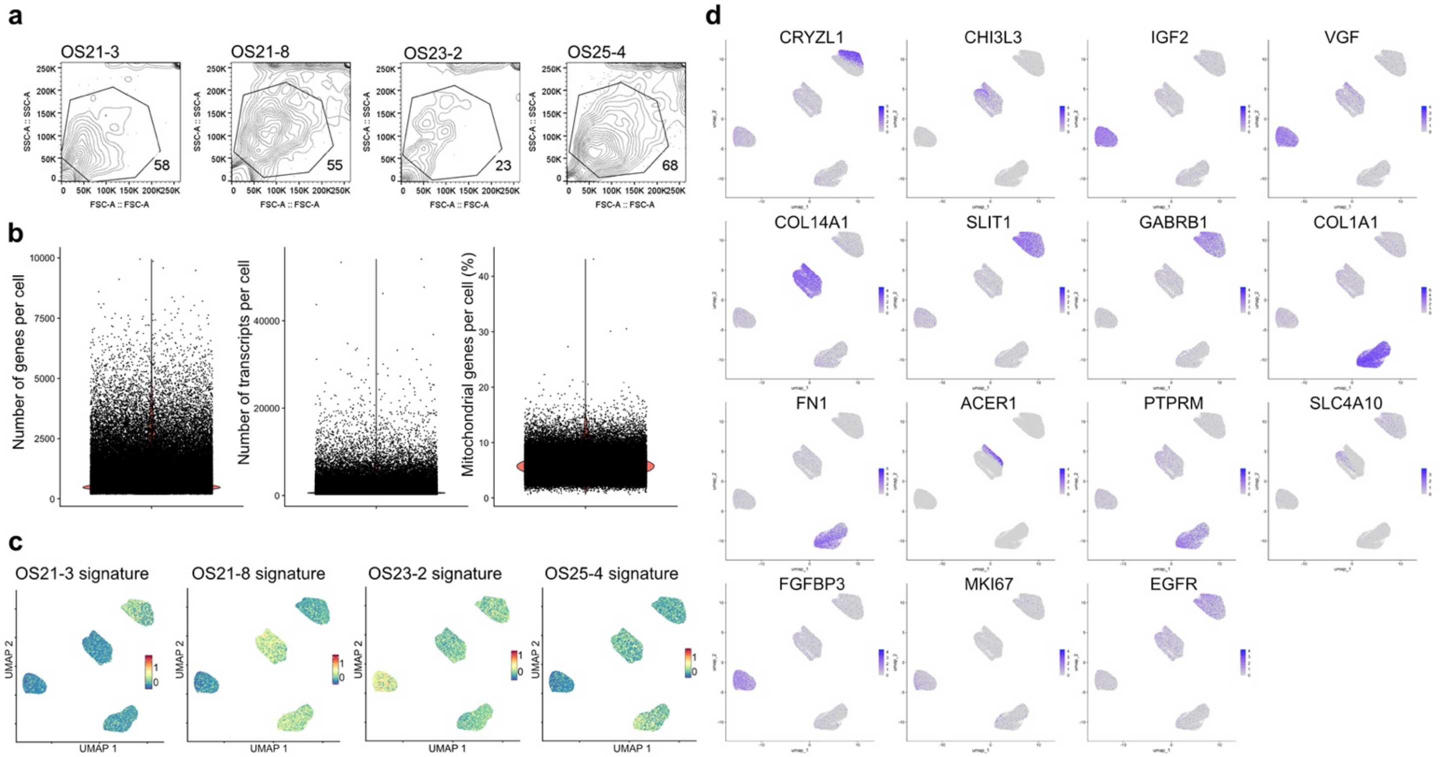


**Supplementary Figure 4.** Quality control for single-cell RNA seq experiment. **a**) Gating of primary GBM neurosphere cells before co-culture to determine cell sizes. **b**) Quality control of single cell RNA seq data. Left panel – number of genes detected in each cell. Middle panel – number of transcript molecules detected in each cell. Right panel – mitochondrial genes percentage per cell. **c**) Confirmation of single cell identity after co-culture and STAR-FACS. Average expression of signatures derived from neurosphere cells is shown. **d**) Single-cell expression of cluster-specific markers, proliferation marker (MKI67) and EGFR.

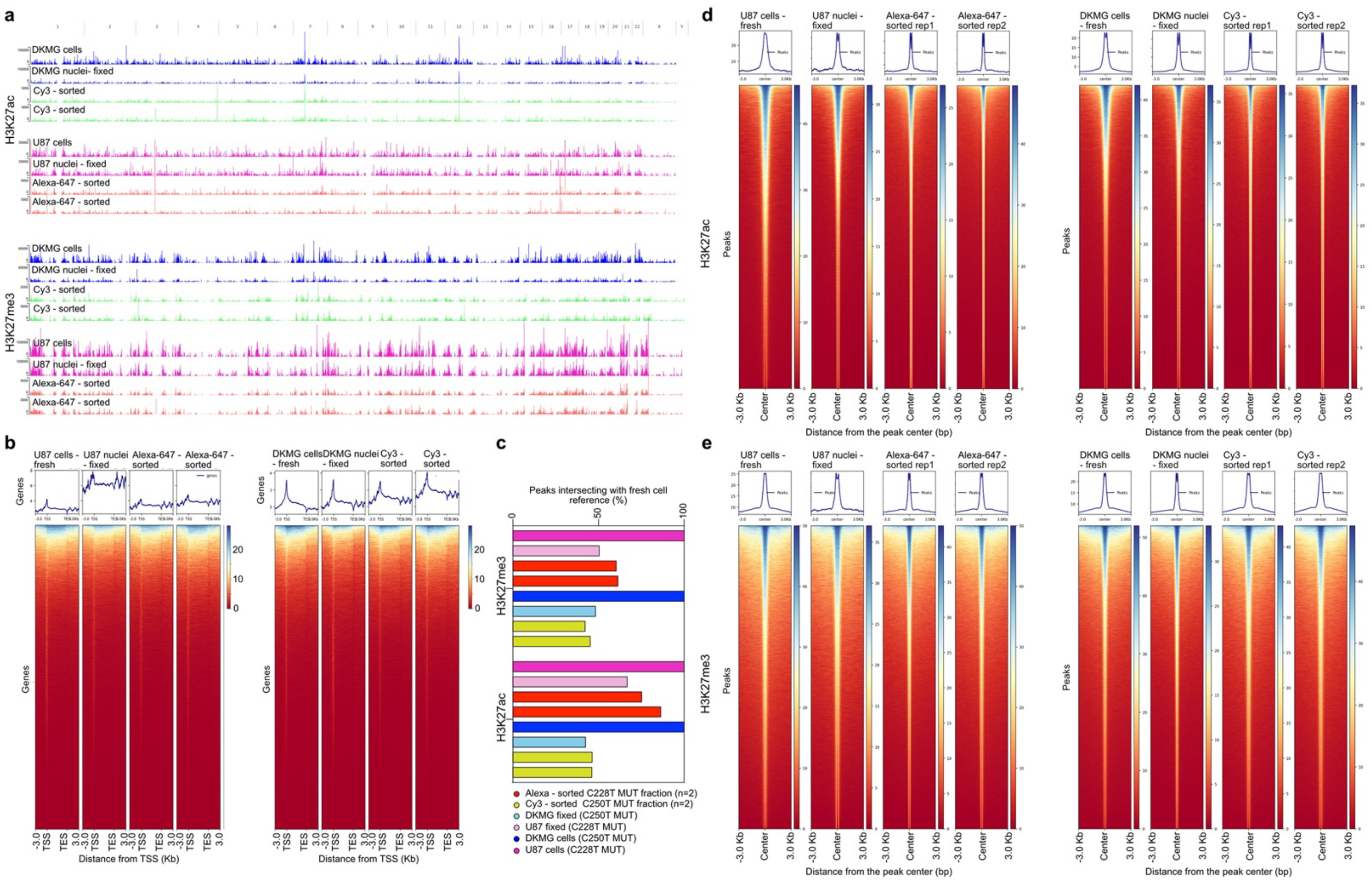


**Supplementary Figure 5.** Histone mark distribution. **a**) Genome-wide distribution of H3K27me3 and H3K27ac profiled by CUT&Tag on fresh cells and nuclei extracted from fixed cells and STAR-FACS sorted fractions from U87-MG and DK-MG mix. Chromosomal locations are indicated at the top. **b**) Distribution of H3K27me3 mark around the transctription start (TSS) and end sites (TES). **c-d**) Average and heatmaps of MACS2 peak summits for CUT&Tag profiling of histone modifications on fresh cells and nuclei extracted from fixed cells and STAR-FACS sorted fractions from U87-MG and DK-MG mix. **c**) Percent intersection of narrow peaks called in each sample. U87-MG and DKMG fresh cell samples were set as references (100%). STAR-FACS sorted samples (2 replicates each) represent similar coverage to their corresponding fixed nuclei sample. **d**) Peak representation in CUT&Tag for H3K27me3. Two replicates of STAR-FACS sorted samples are shown. Alexa-647-sorted samples are TERTp C228T mutant, corresponding to U87-MG cells in the mix. Cy3-sorted samples are TERTp C250T mutant and correspond to DK-MG cells in the mix.
